# DEVELOPMENTAL ULTRASTRUTCTURE MICROSCOPY IN DRUG DISCOVERY FROM DIOSCOREA SPECIES

**DOI:** 10.1101/2023.04.07.535999

**Authors:** Joy Ifunanya Odimegwu, Neelam Singh Sangwan, C. S. Chanotiya, Ogbonnia Steve, Odukoya Olukemi Abiodun

## Abstract

Tubers of wild Dioscorea species (yam) are used as low-cost materials for the synthesis of steroids; cortisone and progesterone using the Marker Degradation chemical route. Decoctions from yam leaves and tubers are also used in ethnomedicine for regulating female fertility and alleviating painful periods and menopausal symptoms. It is now known that yam tuber extracts reduce and inhibit the proliferative action of breast cancer cells and reverse cardiovascular diseases.

Due to current urgent needs and global interest in plants’ biodiversity advancement and drug discovery importance: *Dioscorea composita* and *Dioscorea floribunda* shoots were studied with Scanning Electron Microscopy (SEM) to detect anatomical types of trichomes compared to essential oil production as integrated into the ontogeny of the studied shoots and hydro-distilled oils from the same organs were characterized with Gas Chromatography/Mass Spectrometry (GC/MS).

The SEM study revealed the presence of oil glands, the capitate type on the epidermal layers of parts studied which GC/MS analysis further revealed the components of the oils on the leaves to be Farnesene, Citronellyl acetate, Terpinene, Elemol, Nerolidol, Farnesol and Valerenyl acetate. This is the first time these phytoconstituents are reported in the species. In this light, microscopical methods supported by chromatography contribute to the rapid identification of novel lead compounds contributing to important drug discovery.

## INTRODUCTION

Verpoorte in (2002) stated that Pharmacognosists have mostly been very passionate about introducing new technologies into the Pharmacognostic studies. In the 19th century microscopy was introduced for the quality control of pharmaceutical preparations from plants and it has opened up a broad horizon of discoveries.. Tubers of wild yams are used as low cost materials for synthesis of cortisone and progesterone following the *Marker Degradation* route. Decoctions from yam shoots and tubers are also used in regulating women’s fertility, alleviating painful periods and menopausal symptoms. It is now known that yam tuber extracts inhibit proliferative action of breast cancer cell lines (Odimegwu *et al*; 2019) and reverse cardiovascular diseases in post-menopausal women (Wen, *et al*., 2005). This Plant genus belongs to the family Dioscoreaceae, one of the major members of angiosperms (flowering plants). Plants of the order, Dioscoreales are found in temperate and tropical regions worldwide, with the highest diversity being in the seasonally dry tropics of Central South America (Kirizawa *et al*., 2010), the Caribbean (Raz, 2004), South Africa (Knuth, 1924), and Madagascar (Burkill and Perrier, 1950, Wilkin and Thapyai, 2009). *Dioscorea* is of high value but underutilized (Siqueira, 2011). Its value arises from the huge populations of people who depend on it as a staple food. The economically important yam species include *D*.*alata, D. rotundata* and *D. cayenensis*. The tubers are usually the useful part but in some parts of Africa, the leaves are consumed too. Because of current immediate needs and global keenness in plants’ biodiversity advancement and drug discovery importance; Microscopy has advanced a lot in recent times (Eliceiri, *et al*., 2012) e.g. Fluorescence microscopy is a powerful method to study protein function in its natural habitat, the living cell (Starkuviene and Pepperkok, 2007) with the availability of the green fluorescent protein and its spectral variants very useful in drug discovery. *Dioscorea composita* and *Dioscorea floribunda* fresh leaves (Figure 1A&B) and vines were studied with Stereo microscope (SM) and Scanning Electron Microscopy (SEM) (Figure 4A-F) to detect trichomes anatomical types and essential oil production in the species.

**Figure 1.**
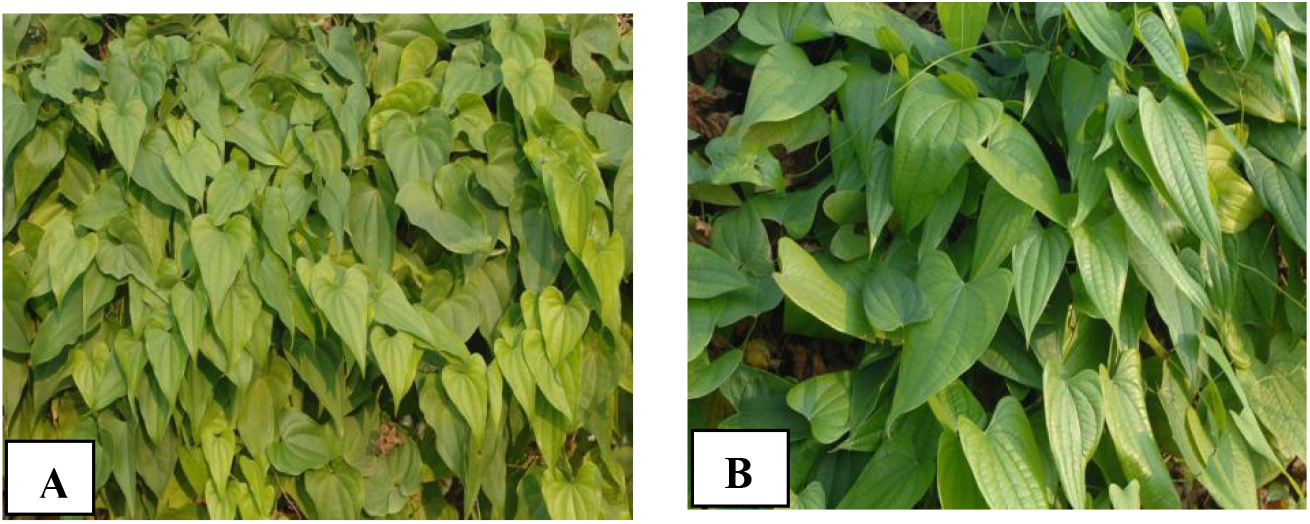
[A] Leaves of *Dioscorea floribunda* [B] Leaves of *Dioscorea Composita*

## METHODOLOGY

Stereo Microscopy (SM) examination of the leaves and shoots were preliminary studies, followed by Scanning Electron Microscopy (SEM) which was used to confirm trichomes anatomical types and hydro-distilled oils from the same organs were characterized with Gas Chromatography/Mass Spectroscopy (GC/MS).

### STEREO MICROSCOPIC STUDIES (SM)

The plant materials were obtained from mature *Dioscorea* plants in the experimental field of the Central Institute of Medicinal and Aromatic Plants (CIMAP) Lucknow, India (26.5oN latitude, 80.5oE longitude, 120m above sea level, subtropical, semi-arid zone with hot summers and cold winters). They were identified by the Institute’s Botanist. Fresh leaves were collected and prepared in the laboratory and subsequently the epidermal layers of the leaves and vines were studied with the aid of a Stereo dissection microscope, model; Leica MZ6 modular with 6.3:1 zoom magnification ranges between 6.3X and 40X for non-destructive 3-D observation.

### SECTION PREPARATION

Transparent nail polish was applied on both sides of detached leaf as a thick patch (at least one square centimetre and allowed to dry completely. The resultant scab was gently peeled from the leaf and subsequently mounted on a microscope slide without staining.

Each leaf (for both species) was divided into three sampling zones (tip, middle, and base) representing equal distances along the length of the leaf blade. Both abaxial and adaxial surfaces were studied. Shape, form and number of trichomes were noted. The leaf impressions were examined under the microscope at different magnifications 10x, 40x, 100x, and 400x and varying directions of illumination was applied (on top, underneath and side) enabling glands and epidermal cells to be easily visualised. Interesting epidermal structures were studied and photographed.

All observed trichomes were counted as they appeared in the microscopic field and the number recorded. Repeat counts from at least three other distinct microscopic fields were taken too. The stomata types observed were also noted.

### SCANNING ELECTRON MICROSCOPY (SEM)

#### REAGENTS AND EQUIPMENT

Critical point dryer, Sputter coater, Specimen collection supplies (tubes, pipettes, dishes, containers), Phosphate buffers; 0.2 M monobasic sodium phosphate, NaH2PO4, and 0.2 M dibasic sodium phosphate, Na_2_HPO_4_, Fixatives, 2.5% glutaraldehyde in phosphate buffer, Methanol, SEM specimen stubs, desiccators, ethanol, Triton X-100, pair of scissors, formalin-alcohol [70%]-Acetic acid 90:5:5, Scanning Electron Microscope Model JEOL-JSM 7500F.

#### SECTION PREPARATION FOR SEM

Leaf and vine samples obtained from field-grown plants and leaves, vine and roots of tissue cultured plantlets were prepared for fixing by immersing all samples in 2.5 % glutaraldehyde in 0.1M phosphate buffer pH 7.4 (with 0.02% Triton X-100) overnight at 4°C. They were washed three times (each after 5min. duration) in 0.1M phosphate buffer pH 7.4. They were dehydrated in ethanol 10min each cycle in 10%, 30%, 50%, 70%, 95% and 100% sequentially and finally left in the 100% solution to ensure total dehydration. The samples were subsequently air dried under a light bulb in the electron microscopy laboratory and finally mounted onto metal stubs with double sided carbon tape. The sputter coater (SciLab, India) was applied automatically and functions by applying a thin layer of gold on the samples to be studied for 10 min.

## RESULTS

The Stereo microscopy observations (Fig 2) revealed capitate glandular trichomes on the epidermal layers from which essential oils were obtained, capitate trichomes can be of various shapes; short, long, with unicellular or pluri-cellular head etc (Liu and Liu, 2012). The trichomes observed (Fig. 3) were short stalked with bulbous heads of 6-12 cells where the essential oils were sequestered. Essential oils with therapeutic effects have been reported from tubers of *D. alata*, (Gramshaw and Osinowo, 1982) and in rhizomes of *D. japonica* (Miyazawa *et al*., 1997) but not in *D. floribunda* and *D. composita*. It was further observed that the glandular trichomes were spread all over the leaf surfaces (abaxial and adaxial) and vines of the test plants with preponderance on the abaxial surface of the species. It was also observed that on epidermal cells of *D*.*composita*, there were more glands at the base of the leaves’ abaxial surface while for *D. floribunda*, there were more at the tip of the leaves’ abaxial surfaces (Table 1). This difference is significant and can be used in taxonomical studies of the species in future. An ideal technique for examining plant surfaces at high resolution is scanning electron microscopy SEM (Pathan *et al*., 2009), ultra-microscopic studies allows for a more detailed study of dehydrated tissues for more accurate reports on the architecture of the epidermal cells.

**Table 1.** Leaf glandular trichomes population from SM of *D*.*composita* and *D. floribunda*

| Observations | *Composita* (number of trichomes) |  |  | *D.floribunda* (number of trichomes) |  |  |
| --- | --- | --- | --- | --- | --- | --- |
|  | Tip | Middle | Base | Tip | Middle | Base |
| Abaxial surface | 13 ± 2.73 | 15 ± 1.53 | 17 ± 0.99 | 8 ± 0.60 | 12 ± 1.84 | 6 ± 1.17 |
| Adaxial surface | 2 ± 0.88 | 2 ± 1.15 | 4 ± 0.58 | 4 ± 0.44 | 3 ± 0.81 | 3 ± 0.33 |

All values are the mean of each group of observations ± standard error of mean. n =3

**Figure 2.**
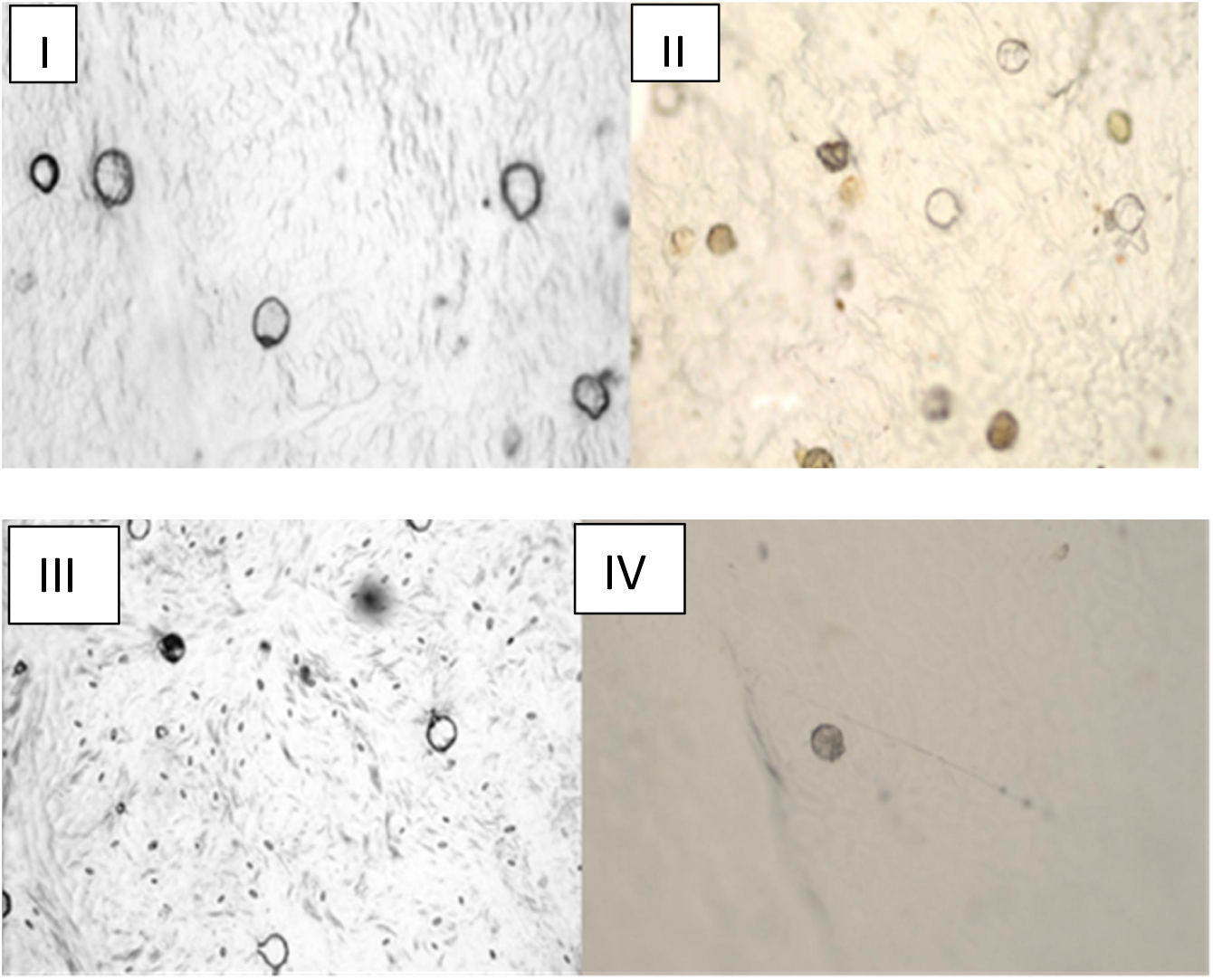
SM image [Mag. X100] of glandular trichomes on *D. floribunda* and *D. floribunda* abaxial and adaxial leaf surfaces (I) *D. floribunda* adaxial view (III) *D. floribunda* adaxial view (II) *D. floribunda* abaxial view of tissue cultured leaf, (IV) *D. floribunda* adaxial view of tissue cultured leaf

**Figure 3.**
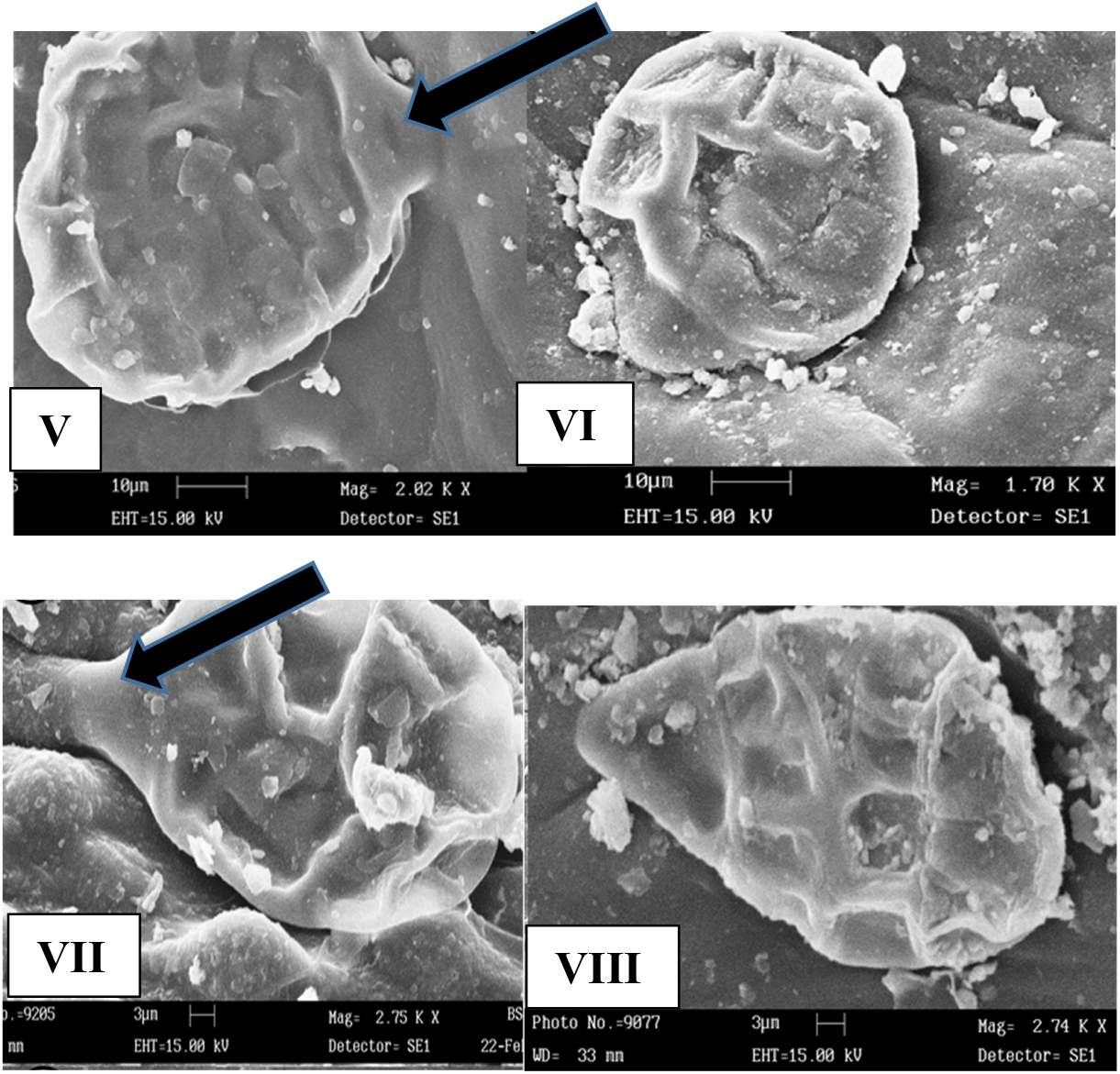
SEM image of glandular trichomes (V). *D. floribunda* leaf surface abaxial view showing point of attachment of glandular trichomes to leaf (black arrow).(VI). *D. floribunda* leaf surface adaxial. (VII and VIII). *D. floribunda* vine surface trichomes.

The SEM study (Fig. 3 & 4) revealed presence of capitate oil glands on the epidermal layers of parts studied, striated lines, radiating from the glandular trichome head were observed on *D. floribunda* leaves’ surfaces (Fig. 3B) but not on *D. composita* leaves. This differentiating feature could be very useful for precise classification of the species as they share many similarities. Calcium oxalate crystals were discovered in *D. floribunda* (Plate 3E). These had been discovered also in *Mentha* according to the work of Turner *et al*., (2000). Oxalate crystals and were observed in the samples are attributed with the function of deterring herbivory due to their itchiness an allergic reaction to the crystals. Chromatographic analysis (Fig. 5Q and R) revealed compounds in the oils to be terpenes; α-terpinene, Nerolidol, Citronellyl acetate, Farnesol, Elemol, α-farnesene, Valerenyl acetate etc. This is the first time these terpenes are reported in the species. In this light, microscopical methods supported by chromatography contribute to rapid identification of novel lead compounds contributing to important drug discovery. The GC-MS analysis revealed essential oils, the most abundant of which was an insecticide, Elemol which were contained in glandular trichomes (a novel finding in *Dioscorea* species) revealed by stereo and scanning electron microscopy.

**Figure 4.**
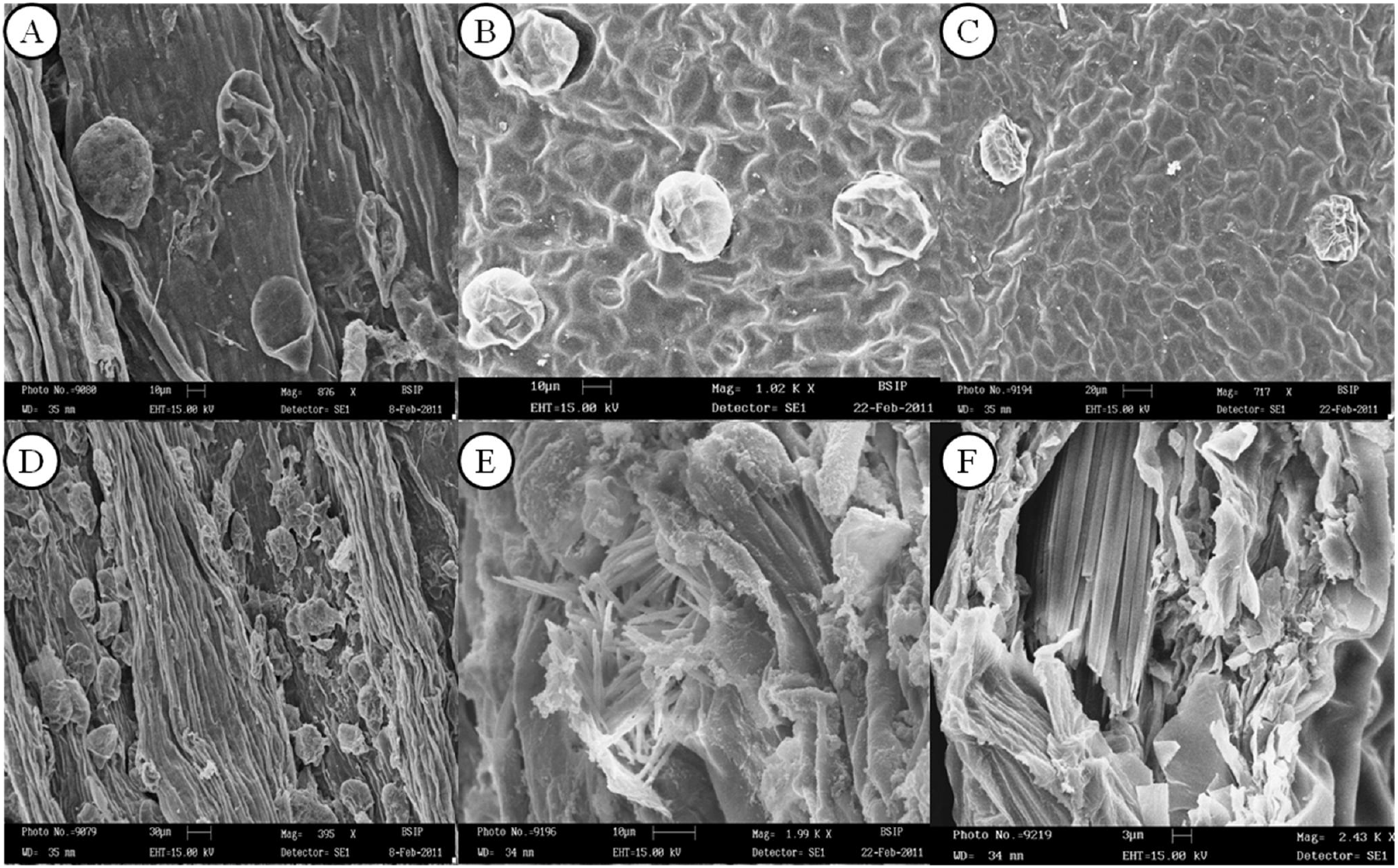
SEM images of *D. composita* and *D. floribunda* leaf, vines and roots’ internal structures. (A) Stem of tissue cultured *D. floribunda* plantlet vine with capitate trichomes (B). Tissue cultured *D. floribunda* leaf surface showing capitate glandular trichomes abaxial and anisocystic stoma, (C). Tissue cultured *D. floribunda* leaf surface showing capitate glandular trichomes adaxial, (D). Stem of tissue cultured *D. floribunda* plantlet showing many capitate glandular trichomes, (E). Tissue cultured *D. floribunda* roots showing oxalate crystals. (F). *D. floribunda* seed transverse section showing neat row of oxalate crystals.

**Figure 5.**
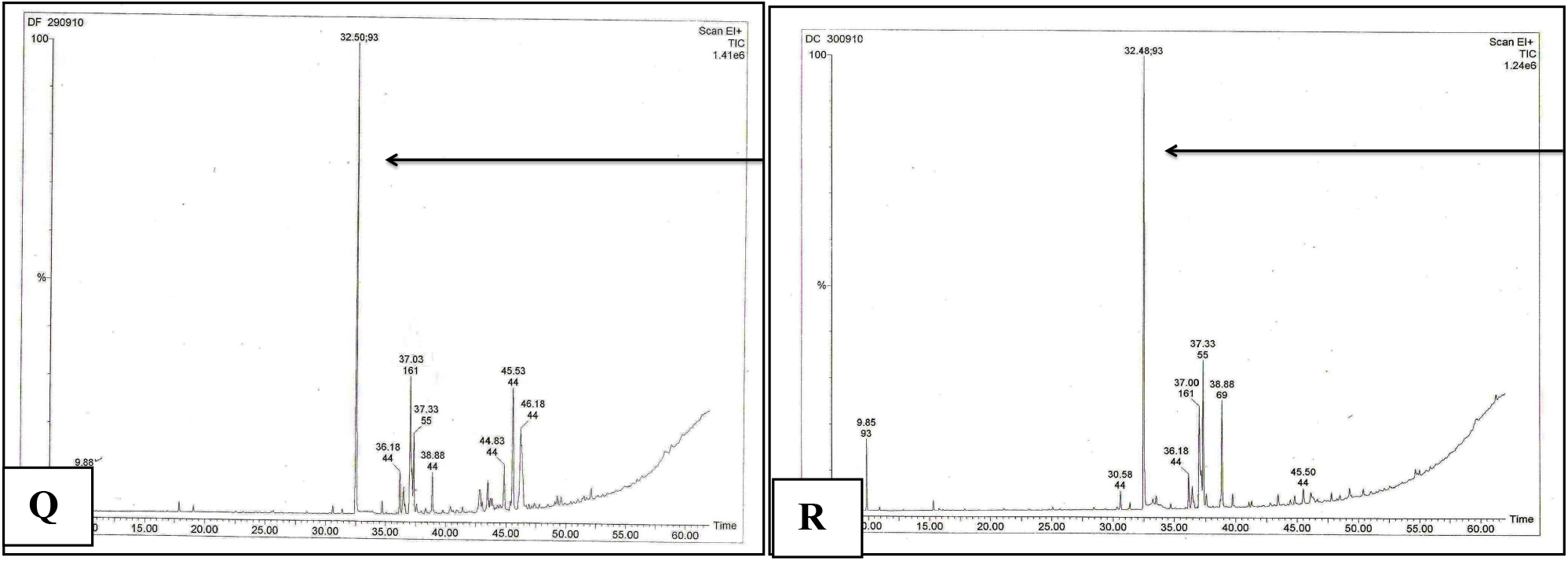
[**Q**] Total ion chromatogram of *D. composita* leaves oil [**R**] Total ion chromatogram of *D. floribunda* leaves oil arrows pointing at Elemol peaks

